# A flexible network of lipid droplet associated proteins support embryonic integrity of *C. elegans*

**DOI:** 10.1101/2022.01.30.478363

**Authors:** Zhe Cao, Chun Wing Fung, Ho Yi Mak

**Affiliations:** Division of Life Science, The Hong Kong University of Science and Technology, Hong Kong SAR, China

**Keywords:** seipin, *seip-1*, lipid droplets, perilipin, Rab18

## Abstract

In addition to coordinating the storage and mobilization of neutral fat, lipid droplets (LDs) are conserved organelles that can accommodate additional cargos in order to support animal development. However, it is unclear if each type of cargo is matched with a specific subset of LDs. Is the identity of each subset rigidly maintained? Here, we report that SEIP-1/seipin defines a subset of oocyte LDs that are required for proper eggshell formation in *C. elegans*. Using a photoconvertible fluorescent protein-based imaging assay, we found that SEIP-1 positive LDs were selectively depleted after fertilization, coincident of the formation of a lipid-rich permeability barrier of the eggshell. Loss of SEIP-1 function caused impenetrant embryonic arrest, which could be suppressed by depleting a LD-structural protein PLIN-1/perilipin. The embryonic development of *seip-1; plin-1* mutant in turn depended on another LD-associated protein, RAB-18/Rab18. Our results are consistent with the hypothesis that SEIP-1 dependent and independent mechanisms act redundantly to ensure the packaging and export of lipid-rich permeability barrier constituents via LDs. The identity of these LDs, as defined by their associated proteins, exhibits unexpected plasticity that ultimately ensures the survival of embryos ex utero.

## Introduction

The deposition of fat is an essential process in the maturation of female germ cells in animals. Such maternal contribution of fat provides energy and membrane precursors that support early embryonic development. Accordingly, lipid droplets (LDs), which are evolutionarily conserved organelles that coordinate fat storage and utilization, are readily detected in vertebrate and invertebrate oocytes and embryos (Welte, 2015; Chen et al., 2020; Ibayashi et al., 2021; Mosquera et al., 2021). The structure of LDs are distinct from other intracellular organelles because a phospholipid monolayer serves as the delimiting membrane (Tauchi-Sato et al., 2002; Walther et al., 2017; Olzmann and Carvalho, 2019). This has led to a model that LDs bud from the outer leaflet of the endoplasmic reticulum (ER) and maintain contact with the ER via protein- and membrane-bridges. The core of LDs contains neutral lipids, such as triacylglycerol (TAG) or cholesterol ester. It is known that the composition of the neutral lipid core varies in a cell type- and nutrient-dependent manner, which in part reflects the demand and supply of specific lipid species (Fu et al., 2014; Molenaar et al., 2021).

Recent evidence suggests that subpopulations of LDs within a single cell can be further distinguished by their association with metabolic enzymes or ER subdomains (Wilfling et al., 2013; Thul et al., 2017; Cao et al., 2019). For example, a subset of LDs in the *C. elegans* intestinal cells associate with a tubular ER-subdomain, defined by the preferential enrichment of the seipin ortholog, SEIP-1 (Cao et al., 2019). Such enrichment is dependent on endogenous polyunsaturated fatty acids and cyclopropane fatty acids that are derived from the bacterial diet, hinting at a link between specific fatty acid availability and LD diversity. In humans, recessive loss-of-function mutations in seipin cause generalized lipodystrophy, which has been attributed to its role in supporting LD biogenesis and expansion (Magré et al., 2001; Payne et al., 2008;Cartwright and Goodman, 2012; Chen et al., 2012; Salo et al., 2016; Wang et al., 2016). In *C. elegans*, the loss of SEIP-1 function reduces the size of a subset of LDs in intestinal cells and perturbs eggshell formation and embryonic development (Cao et al., 2019; Bai et al., 2020). Thus far, it is not completely understood how seipin deficiency at the subcellular level contributes to diverse phenotypes at the organismal level.

The eggshell formation of *C. elegans* is a hierarchical process that demands the sequential secretion of protein- and lipid-rich material into the extracellular space after fertilization (Johnston and Dennis, 2012; Olson et al., 2012; Stein and Golden, 2018). The eggshell protects the embryo from mechanical and osmotic shock to ensure proper development after its expulsion from the mother. Specifically, the lipid-rich layer of the eggshell serves as a permeability barrier that prevents uncontrolled influx of water. Based on genetic analysis, it has been proposed that the permeability barrier is composed of ascarosides, which are sugar-fatty acid conjugates (Olson et al., 2012). However, it is unclear how ascarosides are packaged and exported from the zygote.

In this paper, we investigated the role of SEIP-1 in supporting permeability barrier formation. We were motivated by our observation that *seip-1* mutant worms had an impenetrant embryonic arrest phenotype. Similar to intestinal cells, we observed a subset of oocyte and embryonic LDs that were surrounded by SEIP-1 positive ER. Loss of SEIP-1 function led to the appearance of abnormally large LDs in oocytes. Our attempt to identify genetic modulators of the embryonic arrest phenotype also revealed that the loss of specific LD-associated proteins paradoxically compensated for SEIP-1 deficiency. We propose that multiple ensembles of LD-associated proteins support redundant mechanisms for LDs to accept specialized cargoes. In *C. elegans* oocytes, such SEIP-1 dependent and independent pathways presumably ensure the packaging of ascarosides in LDs prior to their export, which is vital for the construction of the eggshell permeability barrier.

## Results and Discussion

### Loss of *seip-1* function causes impenetrant embryonic arrest

The *seip-1(tm4221)* deletion allele (*seip-1(-)* thereafter) was initially annotated as lethal by the National Bioresource Project (Mitani, 2017). Surprisingly, upon outcrossing with wild type worms, we discovered that *seip-1(-)* worms were fertile (median = 51 live progeny per animal) (column ii, Fig. 1A). The significant reduction in viable progeny was due to a large number of eggs that failed to hatch after being laid. In a separate attempt to study the lipid accumulation of *seip-1(-)* worms, we cultured them in the presence of the vital dye BODIPY on standard nematode growth media (NGM) plates. Although the total number of eggs laid by wild type and *seip-1(-)* worms were comparable (Fig. S1B), BODIPY could not penetrate wild type eggs, while ∼ 60% of *seip-1(-)* eggs were stained (Fig. 1B-D). Interestingly, *seip-1(-)* eggs laid by relatively young adults were more prone to BODIPY staining (Fig. S1C). The strong correlation between BODIPY staining and embryonic arrest suggested that the latter might be caused by a structural deficiency of the eggshell (Fig. S1A). Consequently, the penetration of exogenous material may interfere with embryonic development. Because some *seip-1(-)* embryos develop and hatch as L1 larvae, we therefore conclude that SEIP-1 acts in parallel with additional proteins to ensure embryonic viability. Based on the correlation between aberrant BODIPY staining of *seip-1(-)* embryos and the age of their mothers, it is plausible that the SEIP-1-independent mechanism is triggered at least one day after the initiation of egg laying.

**Figure 1.**
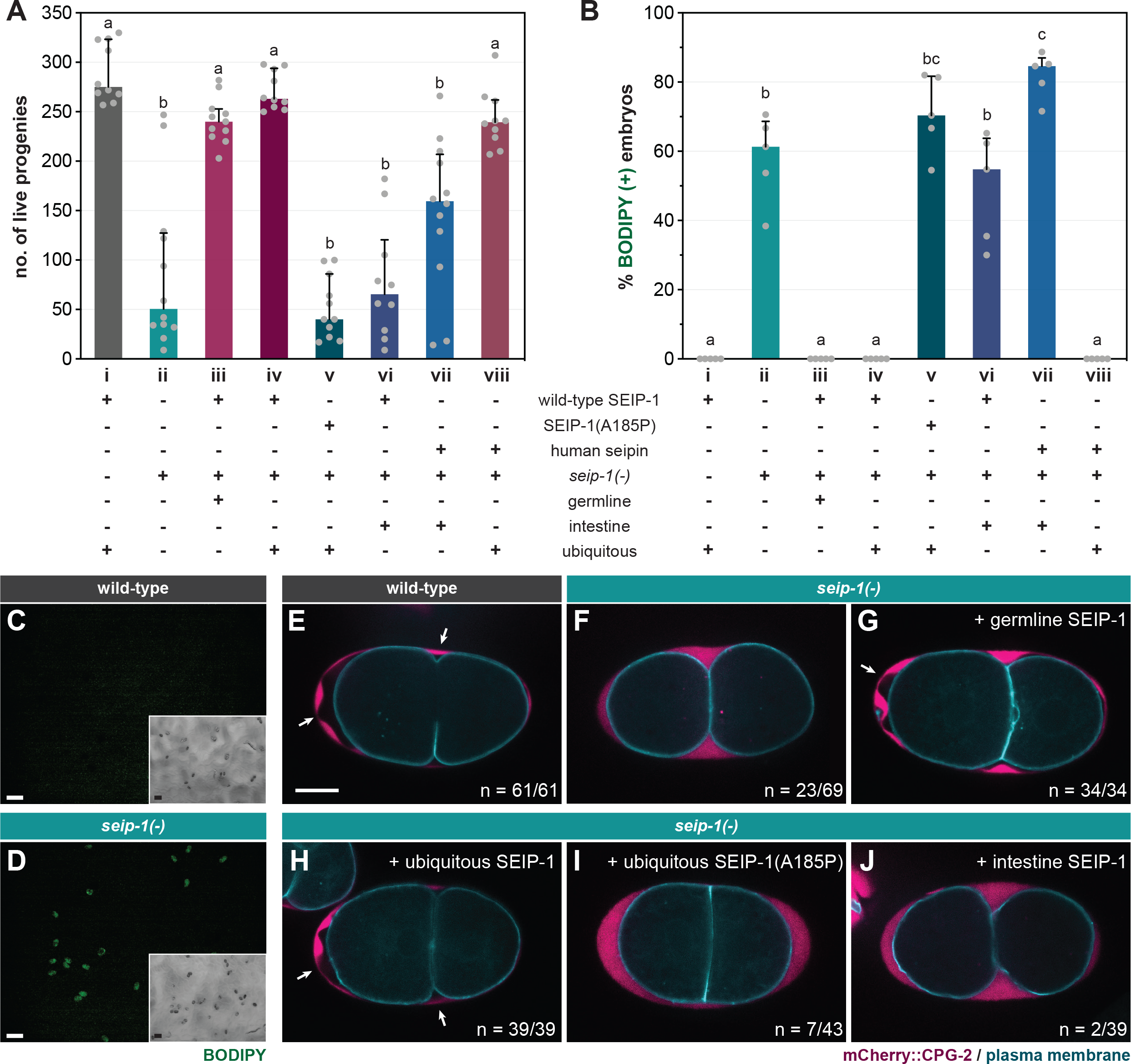
SEIP-1 is required for eggshell integrity of *C. elegans*. (A) The total number of live progeny from individual animals. At least 10 animals of each genotype were scored. Median with interquartile range is displayed (applies to all subsequent bar charts and scatter plots). Groups with different letters are significantly different (ordinary one-way ANOVA with Turkey’s multiple comparisons test, p<0.01). (B) As in (A), but with the percentage of BODIPY-stained embryos quantified in a defined time window. Five independent biological samples were scored, each stemming from four 1-day-old adults. For detailed experimental setup, refer to Methods and Materials. (C) Staining of embryos laid by 1-day-old wild-type (WT) adults with the fluorescent BODIPY 493/503 dye. The dye failed to penetrate WT eggs as shown in the fluorescence image. Inset, the corresponding bright field image. Scale bar = 100μm. (D) As in (C), but with embryos laid by *seip-1(tm4221)* mutants (referred to as *seip-1(-)* in all subsequent figures). The penetration and subsequent accumulation of the BODIPY dye in a subset of embryos is shown. (E) Visualization of the permeability barrier (PB) in a representative 2-cell stage embryo isolated from a 1-day-old WT adult. PB delimits the peri-embryonic space (PES) from the perivitelline space (PVS). White arrows point to PES [nonfluorescent region between PVS, marked by endogenous mCherry::CPG-2 (*hj340*), and the plasma membrane, marked by GFP::PH(PLC1δ1) (*itIs38*)], which is absent if PB formation is impaired. A single focal plane is shown. mCherry and GFP are pseudocolored magenta and cyan, respectively. The schematic layout of the eggshell is shown in Figure S1A. n=number of embryos with PES / total number of embryos examined. Scale bar = 10 μm. (F) As in (E), but with the *seip-1(-)* mutant. (G-J) As in (F), but with the transgenic expression of germline SEIP-1 (*hjSi502*), ubiquitous SEIP-1 (*hjSi189*), ubiquitous SEIP-1(A185P) (*hjSi541*), or intestinal SEIP-1 (*hjSi3*) as indicated.

### A subpopulation of *seip-1(-)* embryos lack the permeability barrier

The structural integrity of the *C. elegans* eggshell is critical for embryonic development, because defects in the eggshell can cause mitotic and polarity defects (Olson et al., 2012). Upon fertilization of the oocyte in the spermatheca, the *C. elegans* eggshell is formed by the hierarchical establishment of four discrete protein- or lipid-rich layers. Starting with the outermost vitellin layer (VL), the chitin layer (CL), the chondroitin proteoglycan layer (CPG), and the permeability barrier were sequentially formed toward the embryonic plasma membrane (Fig. S1A). We noted that the embryonic arrest phenotype of *seip-1(-)* worms was similar to a class of mutants that fail to form the permeability barrier properly. Accordingly, a large number of *perm-1* and *dgtr-1* embryos could be stained when they were laid on BODIPY-containing NGM plates (Fig. S1D). The permeability barrier separates the perivitelline space and the peri-embryonic space (PES) (Fig. S1A). In wild type worms, the permeability barrier retains the chondroitin proteoglycan CPG-2 in the perivitelline space and exclude it from the peri-embryonic space (Fig. 1E). In contrast, we found abnormal accumulation of CPG-2 near the plasma membrane in a fraction of *seip-1(-)* embryos (Fig. 1F), similar to *perm-1* and *dgtr-1* deficient embryos (Olson et al., 2012). Live imaging of wild type embryos *in utero* indicated the emergence of PES upon the cortical ruffling of plasma membrane (Video S1) (Green et al., 2008). PES expands during the pseudocleavage (Goldstein et al., 1993) and is likely finalized following the first mitotic cleavage, which results in two-cell-stage embryos (Video S1). In contrast, PES failed to emerge even after the first mitotic cleavage of some *seip-1(-)* embryos, whereas the exocytosis of CPG-2 was unperturbed (Video S2). Our results are consistent with a recent report (Bai et al., 2020), and implied that SEIP-1 is necessary for the proper formation of the permeability barrier, but not other layers of the eggshell.

### A lipodystrophy-associated mutation at a conserved residue impairs SEIP-1 function

The Alanine 212 to Proline (A212P) mutation in human seipin causes congenital generalized lipodystrophy type 2 disease (CGL2) (Magré et al., 2001). Based on primary sequence alignment, we noted that A185 of SEIP-1 is orthologous to A212 of human seipin (Fig. S2A). To assess if the conserved alanine is important for SEIP-1 function, we generated single-copy transgenes that expressed either SEIP-1(wild-type)::GFP (*hjSi189*) or SEIP-1(A185P)::GFP (*hjSi541*), driven by the ubiquitous *dpy-30* promoter. Wild type SEIP-1::GFP supported the formation of the embryonic permeability barrier (Fig. 1H) and restored the fertility of *seip-1(-)* mutants (Fig. 1A, column iv). Correspondingly, no BODIPY-stained embryos were observed (Fig. 1B, column iv). In contrast, SEIP-1(A185P) failed to suppress the permeability barrier defects of *seip-1(-)* mutant worms (Fig. 1I; Fig. 1A, column v; Fig. 1B, column v). To complement our single-copy transgene strategy, we engineered a knock-in *seip-1(A185P)* allele (*hj158*) using CRISPR. Based on our homology directed repair (HDR) methodology, an “orphan” loxP sequence was inserted between the stop codon and 3’-UTR of *seip-1(A185P)* (Fig. S2B). We therefore constructed a control wild-type *seip-1* allele (*hj156*) with a loxP site inserted at the same position as in *hj158* (Fig. S2B). The fertility of *seip-1(hj156)* animals was comparable to that of wild type. However, the fertility of *seip-1(hj158)* (i.e. A185P) was reduced to a level similar to *seip-1(-)* animals, with a corresponding increase in BODIPY-positive embryos (Fig. S2C-D). We conclude that the A185P mutation disrupts SEIP-1 function, similar to the effect of A212P to human seipin.

### Expression of SEIP-1 in the germline supports embryonic development

Endogenous SEIP-1 is expressed in both the intestine and the germline (Cao et al., 2019; Bai et al., 2020). In *C. elegans*, the proper embryonic development is dependent on the supply of intestine-derived yolk proteins (Grant and Hirsh, 1999; Ezcurra et al., 2018). To this end, we sought to determine the site of SEIP-1 action for proper embryogenesis. Germline-specific expression of SEIP-1 with the *sun-1* promoter (*hjSi502*) fully rescued the defect of *seip-1(-)* embryos (Fig. 1G; Fig. 1B, column iii), thereby restoring the fertility of *seip-1(-)* worms to the wild-type level (Fig. 1A, column iii). In contrast, intestine-specific expression of SEIP-1 with the *vha-6* promoter (*hjSi3*) did not rescue the embryonic defect (Fig. 1J; Fig. 1B, column vi) or the fertility of *seip-1(-)* mutants (Fig. 1A, column vi). Similarly, ubiquitous but not intestine-specific expression of human seipin rescued the defects of *seip-1(-)* embryos (Fig. 1A, columns vii and viii; Fig. 1B, columns vii and viii). Taken together, a conserved function of SEIP-1/seipin is required in the germline to support embryonic development.

### SEIP-1 regulates LD morphology in the germline

We previously demonstrated that SEIP-1 regulates the proper expansion of intestinal LDs (Cao et al., 2019), using the mRuby-tagged *C. elegans* diacylglycerol *O*-acyltransferase 2 ortholog, DGAT-2 (Xu et al., 2012), as the LD marker. Prompted by the germline-specific function of SEIP-1, we examined the morphology of germline LDs in *seip-1(-)* worms. Because DGAT-2 is primarily expressed in differentiated intestinal cells, we used the sole *C. elegans* perilipin ortholog, MDT-28/PLIN-1 (for simplicity, referred to as PLIN-1 in subsequent text), which is ubiquitously expressed (Na et al., 2015), as a germline LD marker. We engineered a knock-in allele (*hj178*) for which the two long isoforms of PLIN-1 (PLIN-1a/c) were tagged with GFP at their C-termini (Fig. 2B). Consistent with our previous finding, when compared to wild-type animals, intestinal PLIN-1-labeled LDs were diminished in size in *seip-1(-)* worms (Fig. 2C-D, inset i). Surprisingly, we found that in both oocytes (inset ii) and embryos (inset iii), abnormally large or small LDs were present in *seip-1(-)* mutants (Fig. 2D), in contrast to the more uniformly sized LDs in the wild-type germline (Fig.2C). As a result, the size range of LDs in *seip-1(-)* germline was larger than that in wild type (Fig. 2E). We made similar observations in *seip-1(A185P)* loss of function mutants (Fig. S2E-G). Such aberrant LD morphology in the SEIP-1 deficient germline is reminiscent of that in yeast and cell line models when seipin orthologs are depleted (Szymanski et al., 2007; Fei et al., 2008; Salo et al., 2016; Wang et al., 2016; Chung et al., 2019). Our results indicate that seipin deficiency causes distinct LD defects in proliferating versus differentiated cells (such as *C. elegans* intestinal cells).

**Figure 2.**
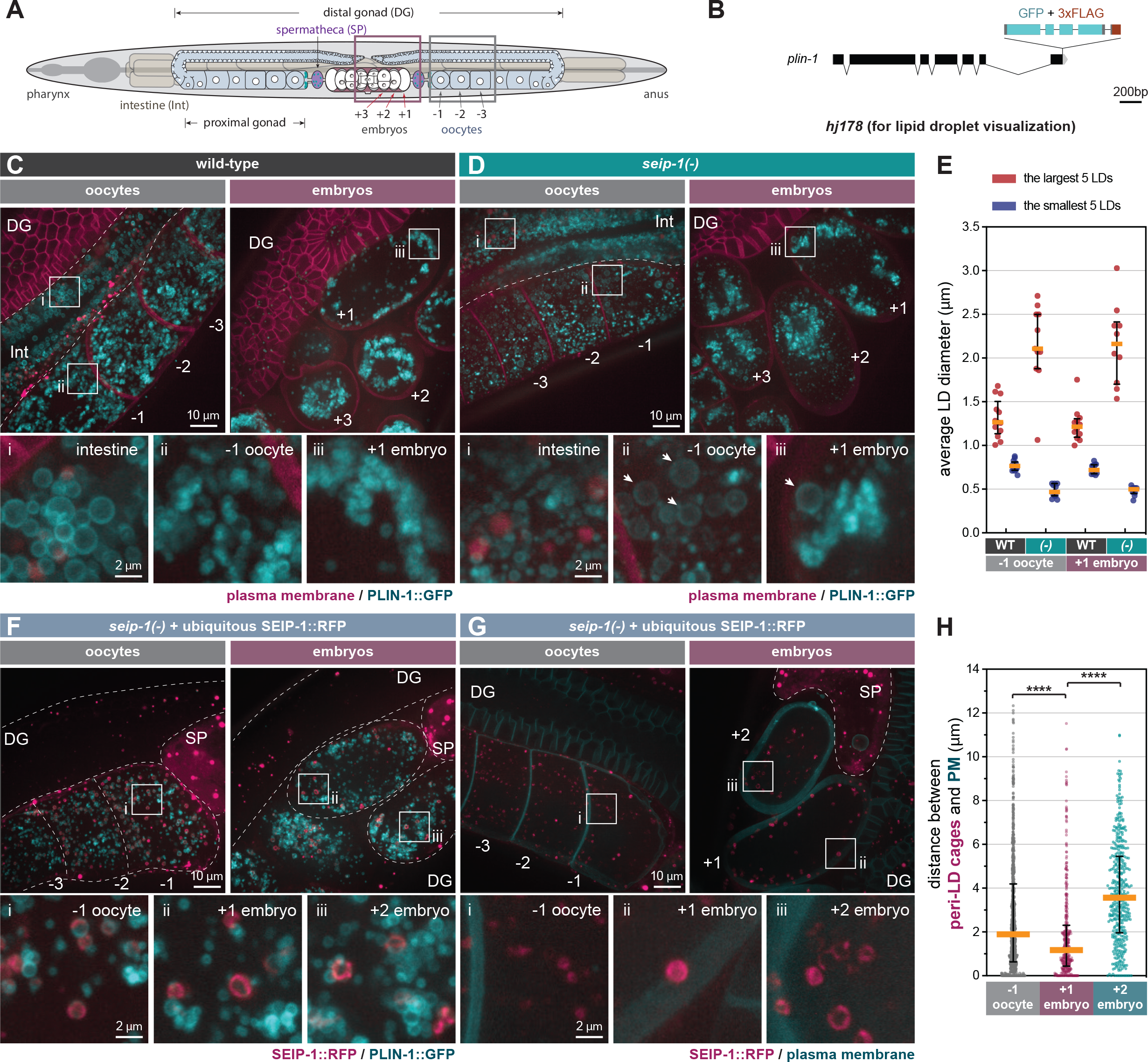
SEIP-1 localizes to peri-lipid droplet (LD) cages and regulates LD morphology in the germline. (A) The anatomy of an adult-stage *C. elegans*. Plum and grey boxes frame the region of interest (ROI) for imaging embryos and oocytes, respectively. (B) Schematic representation of a *plin-1* (*W01A8*.*1*) knock-in allele (*hj178*). Two isoforms of PLIN-1 (PLIN-1a and c), but not PLIN-1b, are fused with GFP at the C-terminus. (C) Visualization of LDs using PLIN-1::GFP (*hj178*) in a 1-day-old wild-type (WT) adult. mCherry::PH(PLC1d1) (*ltIs44*) labels PM in the germline. mCherry and GFP are pseudocolored magenta and cyan, respectively. Dotted lines mark the boundary between different tissues or embryos. Boxed regions were magnified 5x and displayed at the bottom. A projection of 4.5 μm z stack reconstituted from 10 focal planes is shown. For anatomical positions of the ROI, refer to (A). (D) As in (C), but with a *seip-1(-)* mutant. Arrows point to aberrantly enlarged LDs. (E) Average diameter of the largest (crimson) or smallest (navy-blue) five LDs in individual 1-day-old adults. At least 10 animals of each genotype were scored. *(-)* represents *seip-1(-)*. In both oocytes and embryos, when compared to WT, the difference between crimson and navy-blue dots is augmented in *seip-1(-)*. (F) As in (C), but with a 1-day-old adult *seip-1(-)* that expressed SEIP-1::tagRFP (*hjSi434*) and PLIN-1::GFP (*hj178*). The *hjSi434* transgene rescues the embryonic defect in *seip-1(-)* (Figure S3D-E). mCherry and GFP are pseudocolored magenta and cyan, respectively. (G) As in (F), but with a 1-day-old adult *seip-1(-)* that expressed SEIP-1::tagRFP (*hjSi434*) and GFP::PH(PLC1d1) (*itIs38*). (H) The shortest distance between individual SEIP-1::tagRFP labelled peri-LD cages to PM. At least 15 1-day-old *hjSi434; seip-1(-)* adults were scored. Number of peri-LD cages analyzed: -1 oocytes=829; +1 embryos=383; +2 embryos=414. ****p < 0.0001 (ordinary one-way ANOVA with Turkey’s multiple comparisons test).

### Germline SEIP-1 localizes to peri-LD cages

SEIP-1 is enriched at a subdomain of the endoplasmic reticulum (ER), termed the peri-LD cage (Cao et al., 2019), in the *C. elegans* intestine. To investigate the localization of SEIP-1 in the germline, we first examined SEIP-1(WT)::GFP in adult worms that lacked the endogenous SEIP-1 protein. In both oocytes and embryos, SEIP-1(WT)::GFP was targeted to “ring” or “cage”-like structures (Fig. S3A). Similar observations were made in *seip-1(-)* worms that expressed human seipin::GFP ubiquitously (Fig. S3C). In comparison, SEIP-1(A185P)::GFP was rarely found in equivalent structures (Fig. S3B). Such localization defect might be linked to its reduced oligomeric state, as reported for human seipin(A212P) (Binns et al., 2010; Sui et al., 2018). Next, we examined the localization of a SEIP-1::tagRFP fusion protein, relative to the PLIN-1::GFP LD marker. We reasoned that the SEIP-1::tagRFP fusion protein, expressed from a single-copy transgene (*hjSi434*), was functional since it could rescue both the fertility and the osmotic defect of *seip-1(-)* worms (Fig. S3D-E).SEIP-1::tagRFP was enriched in tubular structures around a subset of PLIN-1 positive LDs in the germline (Fig. 2F). Therefore, SEIP-1 appeared to mark a subset of LDs in the germline, similar to our previous observations in the intestine. Additional subcellular compartments, such as cortical granules (Bembenek et al., 2007; Olson et al., 2012) and yolk particles (Sharrock et al.,1990) are known to contribute to early embryogenesis and their reported sizes are similar to that of LDs. Therefore, we asked if SEIP-1 could be found in the proximity of these structures. The cortical granules are exocytosed by canonical anterograde trafficking (Bembenek et al., 2007). Therefore, these granules can be marked with COPII components. We focused on SEC-16A.1, which is one of the two *C. elegans* orthologs of mammalian and yeast SEC16 that is found at ER-exit sites (ERES) and COPII vesicles (Watson et al., 2006). Accordingly, we constructed a knock-in allele that expressed SEC-16A.1 with a C-terminal GFP tag (Fig S3F). For the visualization of yolk particles, we used a published knock-in allele that expressed VIT-2::GFP (Perez et al., 2017). Overall, we did not observe overt colocalization of SEIP-1 and SEC-16A.1 or VIT-2 in oocytes or embryos (Fig. S3G-H). Taken together, we conclude that SEIP-1(+) peri-LD cages are distinct structures in the germline that are necessary for early embryonic development.

### Enrichment of SEIP-1(+) LDs near the plasma membrane upon fertilization

We next sought to understand how SEIP-1(+) LDs might regulate the formation of the permeability barrier. Based on live imaging, we consistently observed LDs and SEIP-1-labeled peri-LD cages near the cell cortex of +1 embryos, but not in -1 oocytes or +2 embryos (Fig. 2A, 2C, 2F-G). To quantify this phenomenon, we compared the localization of SEIP-1 relative to the plasma membrane (Fig. 2G) by measuring the shortest distance from each peri-LD cage to the plasma membrane in cells at either -1, +1, or +2 position. The median distance was reduced from ∼2μm in -1 oocytes and ∼3.8μm in +2 embryos to ∼1μm in +1 embryos (Fig. 2H). In sum, our data is consistent with a model that upon fertilization, SEIP-1(+) LDs are recruited to the cortical region of the zygote to support the formation of permeability barrier. Such distribution pattern of SEIP-1 was not observed in the differentiated intestine (Cao et al., 2019), again highlighting the distinct function of SEIP-1 positive structures in proliferating versus differentiated cells.

### SEIP-1(+) LDs preferentially disappeared during the construction of the permeability barrier

We hypothesized that the redistribution of SEIP-1(+) LDs near the plasma membrane supports the construction of the permeability barrier in newly fertilized embryos. To determine if such redistribution is linked to the catabolism or anabolism of SEIP-1(+) LDs, we developed a photoconversion strategy to “imprint” preexisting SEIP-1 in the oocytes. From the endogenous *seip-1* locus, we expressed the fusion of the monomeric photoconvertible fluorescent protein mKikGR (Habuchi et al., 2008) with SEIP-1 (SEIP-1::mKikGR). In the same worms, we also expressed the ubiquitous LD marker, PLIN-1::GFP. When one-day-old adult worms were exposed to 405nm fluorescence, all green SEIP-1::mKikGR fusion proteins in the germline were converted to red (psuedocolored magenta) (Fig. 3A). These worms were subsequently examined after a one-hour lag. This lag was necessitated when the following events were taken into consideration: ovulation cycle (∼15min) (Huelgas-Morales and Greenstein, 2018), fertilization in the spermatheca (∼5min), and development from fertilization to the first mitotic division (40min) (Stein and Golden, 2018). Therefore, by the time of imaging, the formation of the permeability barrier and the presumptive turnover of oocyte-derived SEIP-1::mKikGR should be complete in +1 embryos. Consistent with our previous study with single-copy transgenes (Cao et al., 2019), endogenous SEIP-1::mKikGR was also targeted to peri-LD cages in the intestine (Fig. 3B, inset i). In the germline, we found fewer pre-existing, photoconverted peri-LD cages in +1 embryos than in -1 oocytes (Fig. 3B, inset ii and iii). Such reduction correlated with a decrease in the photoconverted SEIP-1::mKikGR to PLIN-1::GFP fluorescence intensity ratio (Fig. 3A and 3C).In this case, PLIN-1::GFP fluorescence was used for normalization across different +1 embryos. Altogether, we propose that SEIP-1(+) LDs mobilization and consumption are integral steps of permeability barrier formation.

**Figure 3.**
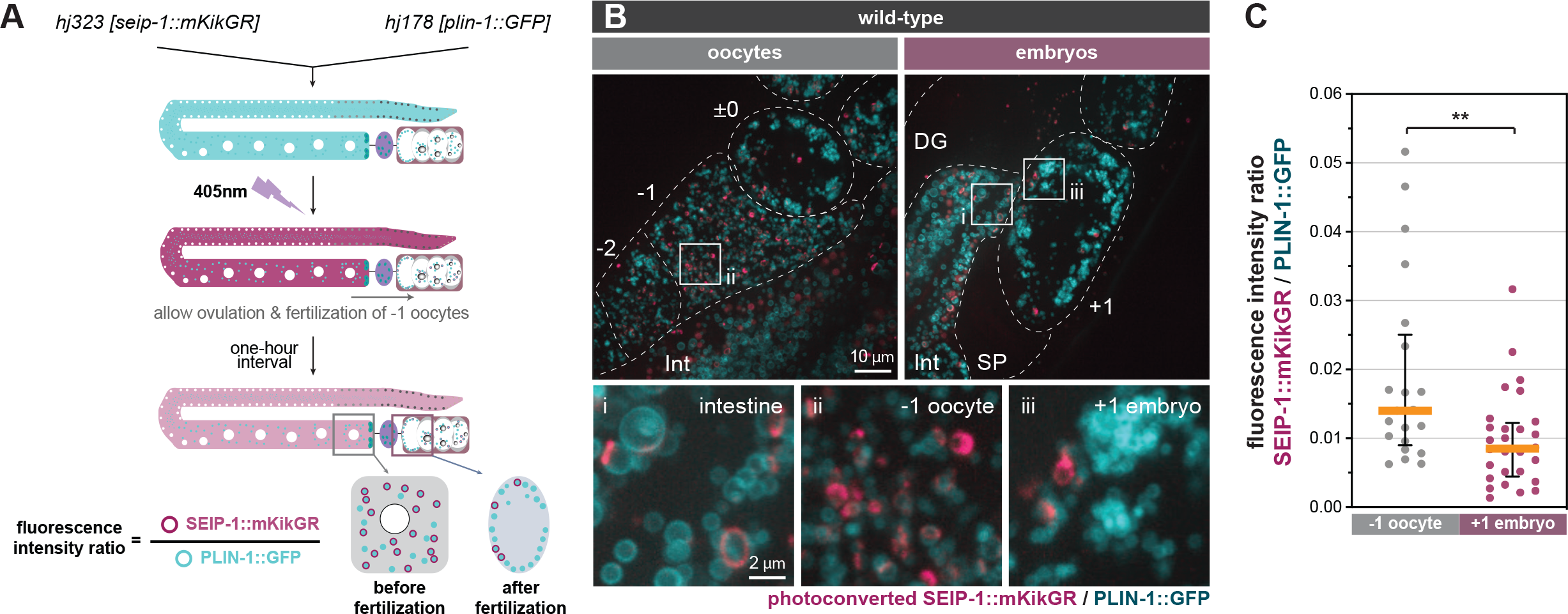
Turnover of SEIP-1 after fertilization. (A) Schematic diagram of the experimental design for the labeling of pre-existing SEIP-1::mKikGR by photoconversion. The initial green (pseudocolored cyan) mKikGR can be photoconverted to red (pseudocolored magenta) by 405nm illumination. The oocytes with photoconverted SEIP-1::mKikGR were allowed to be fertilized in a 1-hour time window prior to imaging. (B) Visualization of photoconverted red SEIP-1::mKikGR (*hj323*) and PLIN-1::GFP (*hj178*). Red mKikGR and GFP are pseudocolored magenta and cyan, respectively. Dotted lines mark the boundary between different tissues or embryos. Boxed regions were magnified 5x and displayed at the bottom. A projection of 4.5μm z stack reconstituted from 10 focal planes is shown. For anatomical positions of the ROI, refer to Figure 2A. (C) Total fluorescence intensity ratio between photoconverted SEIP-1::mKikGR and PLIN-1::GFP in individual 1-day-old adults. Number of animals analyzed: -1 oocytes=21; +1 embryos=28. **p < 0.01 (unpaired *t*-test).

A genetic pathway, consisting of FASN-1, POD-2, PERM-1, and DGTR-1, has previously been shown to control permeability barrier formation (Olson et al., 2012; Stein and Golden, 2018). How does SEIP-1 fit into such pathway? Interesting, the terminal enzyme DGTR-1 is a germline-specific paralog of DGAT-2, which contributes to neutral lipid synthesis in the intestine. We propose that the lipid cargoes, synthesized by DGTR-1, for constructing the permeability barrier are contained in germline LDs. Notably, FASN-1, POD-2, PERM-1 or DGTR-1 deficiency causes embryonic lethality at high penetrance, unlike SEIP-1 deficiency (Fig. 1A-B) (Tagawa et al., 2001; Carvalho et al., 2011). It is plausible that SEIP-1 acts downstream of the cargo synthesis enzymes by specifying a subset of LDs that are destined to be utilized for constructing the permeability barrier.

### Differential requirement for PUFAs in the construction of the permeability barrier

We next sought to understand the molecular basis of incomplete penetrance and high variability of the embryonic defect of *seip-1(-)* worms (Fig. 1A-F, S1C). FAT-3, a polyunsaturated fatty acyl (PUFA) desaturase, is required for targeting SEIP-1 to peri-LD cages in the intestine (Cao et al., 2019). We speculated that the PUFA products of FAT-3 may endow physical properties to ER tubules that are favorable for the recruitment of SEIP-1 to peri-LD cages (Cao et al., 2021). To this end, we asked if FAT-3 deficiency could modify the phenotypes of *seip-1(-)* worms. Indeed, FAT-3 deficiency reduced the fertility of otherwise wild type worms or *seip-1(-)* worms (Fig.4C). Accordingly, *seip-1(-); fat-3(-)* worms displayed a highly penetrant embryonic arrest phenotype, similar to *dgtr-1* or *perm-1* deficient worms. Our data suggests that the products of FAT-3 may partially compensate for SEIP-1 deficiency.

**Figure 4.**
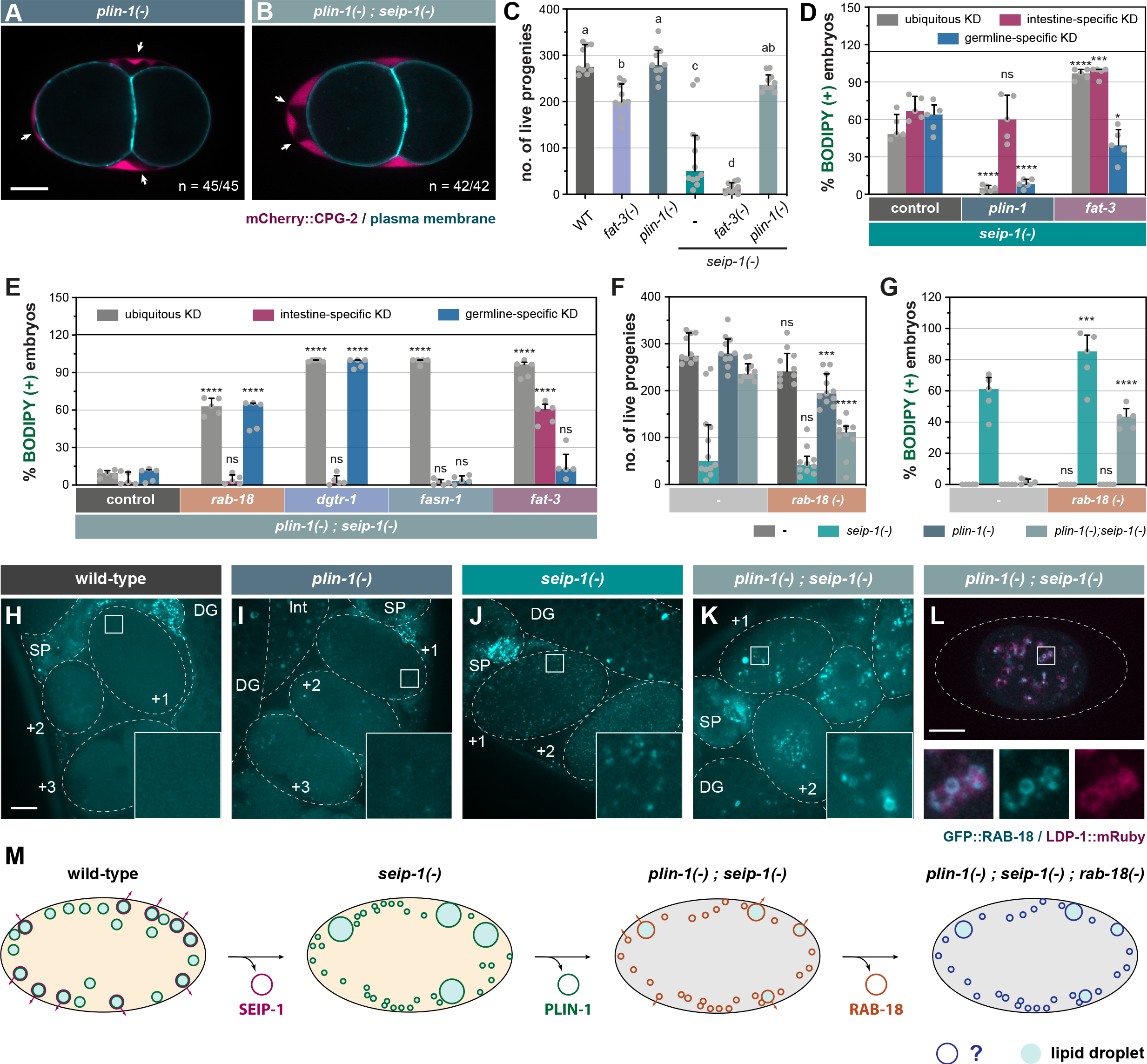
Genetic modifiers of *seip-1(-)* delineate a parallel pathway for eggshell integrity. (A) Visualization of PB in a representative 2-cell stage embryo isolated from a 1-day-old *plin-1(tm1704)* (referred as *plin-1(-)* in all subsequent figures) null adult. White arrows point to PES. A single focal plane is shown. mCherry and GFP are pseudocolored magenta and cyan, respectively. n=number of embryos with PES / total number of embryos examined. Scale bar = 10μm. (B) As in (A), but with *plin-1(-); seip-1(-)* double mutant. (C) The total number of live progeny from individual animals. At least 10 animals of each genotype were scored. Groups with different letters are significantly different (ordinary one-way ANOVA with Turkey’s multiple comparisons test, p<0.01). (D) The percentage of BODIPY-stained embryos with individual genes knocked down in the specified tissues of *seip-1(-)* mutant animals. For every knockdown condition, five independent repeats were scored, each with progeny from four adults. For detailed experimental setup, refer to Methods and Materials. Statistical significance on top of each bar was calculated by comparing each experimental group with its counterpart in the control group (RNAi vertor) (two-way ANOVA with Sidak’s multiple comparisons test). *p < 0.05, ****p < 0.0001. (E) As in (D), but with tissue-specific RNAi in *plin-1(-); seip-1(-)* background as shown. (F) As in (C), but with animals carrying the *rab-18(ok2020)* (hereafter refer to as *rab-18(-)*) allele in the indicated genetic backgrounds. Statistical significance was calculated by comparingeach experimental group (with *rab-18(-)*) with its counterpart in the control group (without *rab-18(-)*) (two-way ANOVA and Sidak’s multiple comparisons test). ns, not significant;***p < 0.001; ****p < 0.0001. (G) As in (F), but with the percentage of BODIPY-stained embryos shown. (H) Visualization of GFP::RAB-18 expressed from its endogenous locus (*hj347*) in an otherwise wild-type 1-day-old adult. Dotted lines mark the boundary between different tissues or embryos. Boxed regions were magnified 5x and shown in the inset. GFP is pseudocolored cyan. (I) As in (G), but in *plin-1(-)* mutant background. (J) As in (G), but in *seip-1(-)* mutant background. (K) As in (G), but in *plin-1(-);seip-1(-)* mutant background. (L) Visualization of GFP::RAB-18 (*hj347*) and LDP-1::mRuby (*hj289*) in an isolated +1 embryo from a 1-day-old *plin-1(-);seip-1(-)* adult. Dotted lines illustrate the boundary of the embryo. LDP-1::mRuby serves as a LD marker. GFP and mRuby are pseudocolored cyan and magenta, respectively. The boxed region was magnified 4x and shown at the bottom. (M) A model on how a flexible network of LD-associated proteins supports embryonic integrity. For all fluorescence images, a projection of 4.5 μm z stack reconstituted from 10 focal planes is shown, scale bar = 10μm.

Next, we expanded our analysis by comprehensively interrogating genes that regulate fatty acid (FA) synthesis and desaturation (Fig. S4A). The percentage of BODIPY-stained embryos (i.e. embryos with defective permeability barrier) was compared after individual genes were depleted by RNAi-based knockdown (KD). In *seip-1(-)* background, ubiquitous depletion of either *fat-3* or *fat-4* increased the number of embryos with permeability barrier defects (Fig. S4B). Accordingly, depleting enzymes upstream of FAT-3/FAT-4 (i.e. FAT-2, FAT-6, ELO-1, and ELO-2) enhanced the embryonic arrest of *seip-1(-)* mutants. Similar effect was not observed when FAT-1 or FAT-5 was depleted (Fig. S4B).

To determine the tissues from which fatty acid desaturases modulated the phenotypes of *seip-1(-)* worms, we repeated our experiments with strains that restricted RNAi to the intestine or germline (Cao et al., 2019; Zou et al., 2019). In *seip-1(-)* mutant background, intestine-specific depletion, but not germline-specific depletion, of FAT-2, FAT-3, or FAT-4 enhanced the embryonic arrest phenotype (Fig. 4D, S4D). Therefore, PUFAs synthesized in the intestine appear to support a mechanism that partially compensates for SEIP-1 deficiency in the germline. We also investigated the dependence on fatty acid desaturases for embryonic development in otherwise wild-type worms. Interestingly, embryonic arrest was observed only when FAT-2 was depleted ubiquitously (Fig. S4C). However, neither intestine-nor germline-specific knockdown of *fat-2* recapitulated phenotypes observed when *fat-2* was knocked down ubiquitously. Our results imply that FAT-2 products made in one tissue can diffuse to other tissues to compensate for local FAT-2 deficiency. Based on the differential requirement of fatty acid desaturases in wild type and *seip-1(-)* worms, multiple mechanisms that demand different PUFAs appear to act redundantly, in order to ensure the formation of the permeability barrier, which is vital for embryonic development.

### PLIN-1 deficiency rescues the embryonic defect of *seip-1(-)* mutants

We inadvertently discovered that PLIN-1 inhibited an alternative mechanism that supported permeability barrier formation in *seip-1(-)* worms. In an attempt to tag all isoforms of PLIN-1 with the mRuby red fluorescent protein, we inserted the mRuby coding sequence to the 5’ end of the *plin-1* gene by CRISPR (Fig. S5A). When we introduced the resultant allele, *hj249*, into *seip-1(-)* worms, we were surprised that the fertility of *plin-1(hj249); seip-1(-)* worms was similar to that of wild type worms (Fig. S5B). Accordingly, the percentage of BODIPY-positive embryos was significantly reduced (Fig. S5C). These phenotypes were not observed in *plin-1(hj178)* when PLIN-1A and PLIN-1C isoforms were fused at their C-terminus to GFP (Fig. S5B-C). Such discrepancy led us to speculate that the mRuby fusion to the N-terminus of PLIN-1 compromised its function, and that *hj249* was a hypomorphic *plin-1* allele that rescued SEIP-1 deficient embryos. To prove our hypothesis, we analyzed worms that carried the *plin-1(tm1704)* deletion allele (Xie et al., 2019) (referred to as *plin-1(-)* hereafter). The *plin-1(-)* worms showed normal permeability barrier formation and fertility (Fig. 4A and C). Similar to *plin-1(hj249); seip-1(-)* worms, the fertility of *plin-1(-); seip-1(-)* worms was comparable to that of wild type worms, and almost all embryos had an intact permeability barrier as indicated by the lack of BODIPY staining (Fig. 4B-C). In addition, the *plin-1(-)* allele rescued the embryonic defects of *seip-1(A185P)* mutants (Fig. S2C-D), suggesting that the suppression by *plin-1(-)* was not restricted to specific *seip-1* loss of function alleles.

Next, we used RNAi to knockdown *plin-1* constitutively, or tissue-specifically in the intestine or germline. We found that constitutive or germline-specific RNAi against *plin-1* suppressed *seip-1(-)* (Fig. 4D). However, intestine-specific RNAi against *plin-1* did not show similar suppression (Fig. 4D). As a complementary approach, we expressed *plin-1* in the germline of *plin-1(-); seip-1(-)* worms, with a *sun-1* promoter driven single-copy transgene. These worms showed embryonic defects similar to *seip-1(-)* worms (Fig. S5B-C). Thus, our results indicate that the loss of PLIN-1 function in the germline is sufficient to bypass the requirement on SEIP-1 for permeability barrier formation. Interestingly, constitutive knockdown of *fat-2* or *fat-3* and intestine-specific knockdown of *fat-3* by RNAi reversed the suppression of *seip-1(-)* associated embryonic defects by *plin-1(-)* (Fig. 4E, S4E). Therefore, *plin-1(-); seip-1(-)* worms appeared to rely on FAT-2 or FAT-3, acting in distinct tissues, to support permeability barrier formation. Our results support the notion that local and diffusible PUFAs are differentially required for embryonic development of *C. elegans*.

### RAB-18 is required for the development of *seip-1(-); plin-1(-)* embryos

Next, we sought to identify proteins that acted redundantly with SEIP-1 to ensure embryonic development. To this end, we inactivated genes encoding known or putative LD-associated proteins by RNAi. We found that constitutive or germline-specific *rab-18* knockdown yielded BODIPY-positive, permeability barrier defective *plin-1(-); seip-1(-)* embryos (Fig. 4E). Intestine-specific RNAi against *rab-18* had no effect (Fig. 4E). Similar observations were made when *dgtr-1*, a gene known to be essential for permeability barrier formation (Olson et al., 2012), was knocked down in *plin-1(-); seip-1(-)* embryos (Fig. 4E). Our results suggest that a RAB-18 acts in the germline, in the absence of SEIP-1, to support permeability barrier formation in a DGTR-1-dependent manner. It should be noted that loss of *dgtr-1* function caused highly penetrant embryonic arrest of otherwise wild type worms (Carvalho et al., 2011). In contrast, worms harboring the *rab-18(ok2020)* (referred to as *rab-18(-)* hereafter) loss of function allele did not arrest as embryos and have comparable brood size as wild type worms (Fig. 4F). Although *rab-18(-)* enhanced the embryonic arrest phenotype of *seip-1(-)* worms by ∼20% (Fig. 4G), there was no significant impact on the brood size (Fig. 4F). The loss of *rab-18* also reduced the brood size of *plin-1(-)* worms by ∼20% (Fig. 4F), but it could not be attributed to embryonic arrest (Fig. 4G). Finally, *rab-18(-)* significantly increased the percentage of arrested embryos from *plin-1(-); seip-1(-)* worms and reduced their brood size accordingly (Fig. 4F-G). Therefore, RAB-18 was specifically required in *plin-1(-); seip-1(-)* worms to support embryonic development.

To elucidate the mechanism by which RAB-18 supported permeability barrier formation in the absence of SEIP-1 and PLIN-1, we investigated the localization of GFP::RAB-18 fusion protein, expressed from its endogenous locus (Fig. S6). It did not localize to distinct structures in wild type and *plin-1(-)* worms (Fig. 4H-I). In *seip-1(-)* worms, GFP::RAB-18 appeared in numerous diffraction-limited puncta throughout the cytoplasm of newly fertilized (+1, +2) embryos (Fig. 4J). Finally, in *plin-1(-); seip-1(-)* embryos, GFP::RAB-18 was found on the LD surface as confirmed by the LDP-1::mRuby marker (Na et al., 2015) (Fig. 4K-L). Interestingly, Rab18 is enriched in ‘immature’ LDs in seipin deficient mammalian cells (Salo et al., 2019). Furthermore, ADRP/Perilipin 2 and Rab18 appeared to compete for LD surface association in cultured mammalian cells (Ozeki et al., 2005). Therefore, it was plausible that the loss of PLIN-1 in *seip-1(-)* embryos similarly permitted the association of RAB-18 with ‘mature’ LDs that contained the cargoes for permeability barrier construction. Taken together, our results support the notion that RAB-18 positive LDs assume an identity that is similar to SEIP-1(+) LDs in otherwise wild type embryos. The association of RAB-18 with LDs is modulated by other resident LD proteins, such as PLIN-1.

### Concluding remarks

In this paper, we used genetic and imaging approaches to reveal a requirement for SEIP-1 in the formation of the permeability barrier, which is part of the *C. elegans* eggshell. Our results suggest that SEIP-1(+) LDs contribute to the packaging and release of lipid-rich ascarosides that are eventually exported to the embryonic extracellular space. To this end, the ability of SEIP-1 to mark a subset of mature LDs seems to be separable from its other established function in supporting the emergence of nascent LDs from the ER (Chung et al., 2019; Salo et al., 2019; Prasanna et al., 2021; Zoni et al., 2021). It is plausible that SEIP-1 enrichment to ER subdomains promotes the assembly of enzymes that are required for ascarosides synthesis, prior to their deposition into specialized LDs. Future efforts will be needed to determine the localization of germline factors that are known to contribute to the permeability barrier formation. It is well-established that mammary epithelial cells export milk fat in the form of LDs (Masedunskas et al., 2017). Interestingly, seipin knockout mice are defective in milk production and lactation (El Zowalaty et al., 2018). Therefore, the involvement of seipin-positive LDs in the export of lipophilic molecules may be a conserved phenomenon. Our discovery that PLIN-1 deficiency can suppress the *seip-1(-)* embryonic arrest phenotype suggests that parallel pathways exist to permit ‘alternative’ LDs to accommodate and release ascarosides (Fig. 4M). Taken together, we propose that the plasticity of the LD surface coat supports a safety mechanism that ensures the construction of the permeability barrier, which is crucial for *C. elegans* eggshell integrity and embryo survival ex utero.

## Methods and Materials

### Strains and transgenes

Bristol N2 was used as the wild-type *C. elegans* strain. All animals were maintained and investigated at 20°C. The following alleles and transgenes were obtained from the Caenorhabditis Genetics Center (CGC): *LG III, unc-119(ed3), rab-18(ok2020), itIs38[(pAA1)pie-1p::GFP::PH(PLC1delta1)+unc-119(+)]; LG IV, fat-3(ok1126); LG V, sid-1(qt78), ltIs44[(pAA173)pie-1p::mCherry::PH(PLC1delta1)+unc-119 (+)]; LG X, vit-2(crg9070). seip-1(tm4221) V* and *plin-1(tm1704) I* alleles were obtained from Dr. Shohei Mitani (National Bioresource Project for the nematode). *rde-1 (mkc36) V* and *mKcSi13 [sun-1p::rde-1::sun-1 3′UTR] II* are gifts from Dr. Di Chen (Model Animal Research Center, Nanjing University). The following single-copy transgenes were used: *hjSi3[vha-6p::seip-1 cDNA::GFP_TEV_3x-FLAG::let-858 3′UTR] II, hjSi189[dpy-30p::seip-1 cDNA::GFP_TEV_3xFLAG::tbb-2 3′UTR] II, hjSi206 [vha-6p::human seipin isoform 2 cDNA(codon optimized)::GFP_TEV_3xFLAG::let-858 3′UTR] II, hjSi223[dpy-30p::human seipin isoform 2 cDNA (codon-optimized)::GFP_TEV_3xFLAG::tbb-2 3′UTR] II, hjSi434[dpy-30p::seip-1 cDNA::tagRFP::tbb-2 3′UTR] II, hjSi494[vha-6p::sid-1 cDNA::dhs-28 3′UTR] I, hjSi502[sun-1p::seip-1 cDNA::GFP_TEV_3xFLAG::sun-1 3′UTR] II, hjSi541[dyp-30p::seip-1(A185P) cDNA::GFP_TEV_3xFLAG::tbb-2 3′UTR] II, hjSi552[sun-1p::plin-1 gDNA::GFP_TEV_3xFLAG::sun-1 3′UTR] II*. The following CRISPR/Cas9-generated alleles were used: *hj158[seip-1(A185P)::stop codon_loxP_seip-1 3’-UTR] V, hj178[plin-1a/c::GFP_TEV_3xFLAG] I, hj249[mRuby_TEV_3xFLAG::plin-1] I, hj289[ldp-1::mRuby_TEV_3xHA] V, hj323[seip-1::mKikGR_SEC_3xFLAG] V, hj340[cpg-2 signal peptide::mCherry_TEV_3xFLAG::cpg-2 w/o signal peptide] III, hj347[GFP_TEV_3xFLAG::rab-18b] III*.

### RNA interference-based knockdown in *C. elegans*

RNA interference (RNAi) was performed by on-plate feeding according to published methods (Neve et al., 2020). The targeting sequence in each RNAi vector was either taken from the Ahringer library or constructed with primers detailed in supplementary table 1. The plasmids were transformed into OP50 [rnc14::DTn10 laczgA::T7pol camFRT] (Neve et al., 2020). Fresh overnight cultures of OP50 RNAi clones were seeded on NGM plates with 0.4 mM IPTG and 100 μg/mL Ampicillin. The seeded plates were stored in the dark and incubated at room temperature for one day prior to experiments. The intestine-specific and germline-specific knockdown was performed as previously described (Cao et al., 2019; Zou et al., 2019). In brief, the single-copy intestine-specific (*hjSi494*) or germline-specific (*mKcSi13*) transgene is used to rescue *sid-1(qt78)* or *rde-1(mKc36)* mutant animals, respectively. The *seip-1(tm4221)* and/or *plin-1(tm1704)* allele were introduced into *hjSi494; sid-1 (qt78)* and *mKcSi13; rde-1(mKc36)* by genetic crosses.

### Fluorescence imaging of *C. elegans*

For all fluorescence imaging, one-day-old adult animals were examined by a spinning disk confocal microscope (AxioObeserver Z1, Carl Zeiss) with a ×5 (numerical aperture (NA) .16) or ×63 (numerical aperture (NA) 1.4 oil) Alpha-Plan-Apochromat objective. Image stacks were acquired with a Neo sCMOS camera (Andor) and a piezo Z stage controlled by the iQ3 software (Andor). For GFP, a 488 nm laser was used for excitation and emitted signals were collected by a 500–550-nm filter. For mCherry, mRuby, tagRFP, and photo-converted mKikGR, a 561 nm laser was used for excitation and emitted signals were collected by a 580.5–653.5-nm filter. Optical sections of images at 0.5μm intervals were exported to Imaris 8 (Bitplane) for processing and 3D reconstruction.

For imaging the germline and embryos of live *C. elegans*, fresh 8% agarose pads were prepared on top of microscope glass slides. 2-3μl of 1.25% (w/v) polystyrene microspheres (Polybead Microspheres 0.05μm, Polysciences) and 0.2mM levamisole (Sigma) in 1×PBS were dropped at the center of the agarose pads for immobilization of adult animals. For imaging the permeability barrier, on 22mm × 22mm cover slips, embryos were dissected from the uterus of one-day-old adults (24h after the mid-L4 larval stage) in 0.8×egg buffer with 0.2mM levamisole in spherical micro-chambers bordered by vaseline. The microscope glass slides were subsequently applied with care to avoid the compression of the embryos.

For photoconversion of mKikGR, individual plates of one-day-old *seip-1(hj323)* adults (24h after the mid-L4 larval stage) were exposed to 400-440nm fluorescence filtered by SZX2-FBV (Olympus) for 30min on a stereomicroscope (SZX16, Olympus) equipped with a LED light source (EXFO). Worms were allowed to rest for 1h at 20°C so that -1 oocytes were ovulated and fertilized prior to confocal microscopy.

### Analysis of fluorescence images

The diameter of LDs present in -1 oocyte and +1 embryos was manually fitted using the spot function in Imaris. The shortest distance between peri-LD cages and plasma membrane was computed by the spot and surface functions of Imaris XT.

### Measurement of total number of live progenies

Animals carrying the *seip-1(tm4221)* allele were backcrossed four times with wild-type males right before measurement. L4 larval P0 animals of each genotype were singled to individual plates and transferred to new plates every 24-48h until egg-laying ceased. Hatched F1 animals were monitored and counted 24-48h after the removal of the P0.

### BODIPY-staining of embryos

Four one-day-old adults (24h after the mid-L4 larval stage) per replicate per strain were transferred to a new plate and allowed to lay eggs at 20°C. After three hours, adults were removed and the plate was stained with 500μl 1mM BODIPY in 1×PBS for 3h. The percentage of BODIPY-positive embryos was calculated using a fluorescence stereomicroscope. For comparison upon knockdown of different genes, the setup was similar as described above except that the age of adults varied in order to ensure both the efficiency and specificity of the knockdown. For constitutive knockdown, L4 larval P0 animals were plated and one-day-old F1 animals were used for scoring the percentage of BODIPY-positive embryos. For intestine-specific knockdown, L4 larval animals were plated and scored 48h later. For germline-specific knockdown, L4 larval animals were plated and scored 24h later.

### Generation of knock-in/knock-out alleles in *C. elegans*

Tagging of the endogenous genes was performed as described previously (Dickinson et al., 2015). In brief, single guide RNAs (sgRNAs, detailed in supplementary table 2) were designed using CHOPCHOP (https://chopchop.rc.fas.harvard.edu/) and cloned into pDD162 (Addgene). The sgRNA/Cas9 vectors were injected into adult *C. elegans* along with the corresponding repair templates and co-injection markers. The transgenic P0s were singled and their progeny were selected with 5mg/mL hygromycin B (InvivoGen). Viable F2s were further screened using a fluorescence stereomicroscope and PCR-based genotyping to verify knock-in alleles. For knocking out a target gene, 2-4 sgRNAs were injected along with co-injection markers. F1s from the transgenic P0s were singled and genotyped for deletion. For each knock-in/knock-out strain, at least two independent alleles were acquired and examined for consistency. All constructed alleles were backcrossed two times with wild-type males before further characterization.

## Acknowledgements

We thank Yan Hao and Joshua Fung for preliminary data. Di Chen, Meng Wang and King L. Chow for RNAi-related strains and reagents. Jihong Bai, Christian Frøkjær-Jensen and Bob Goldstein for Mos1 and CRISPR reagents. Some strains were provided by the CGC, which is funded by NIH Office of Research Infrastructure Programs (P40 OD010440). The work was supported by RGC GRF 16102118 and 16101820 to HYM.

---

**Figure S1.**
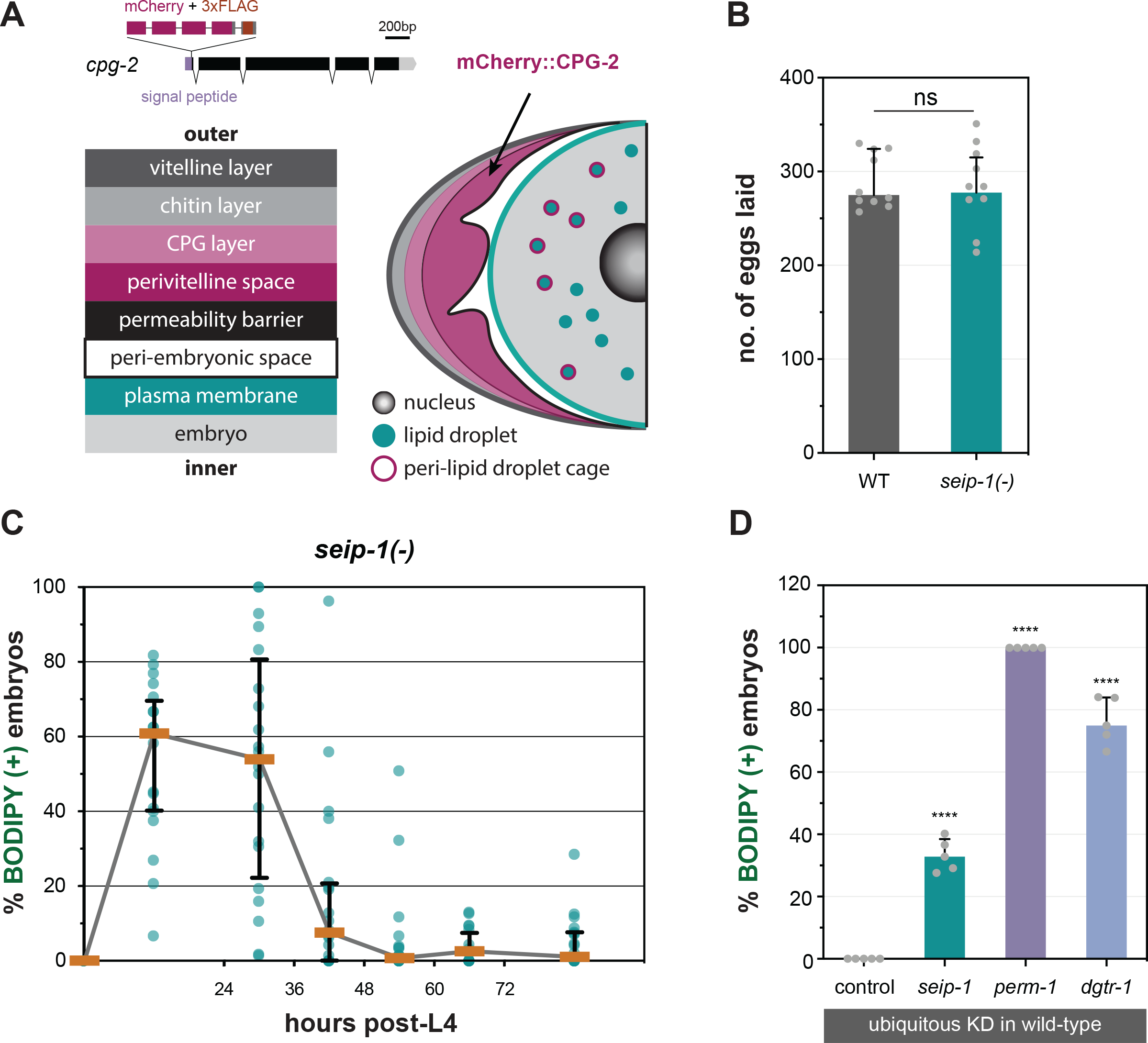
SEIP-1 supports eggshell integrity of *C. elegans*. (A) A schematic diagram summarizing the eggshell composition of wild-type *C. elegans*. The design of the *mCherry::3xFLAG::cpg-2* knockin allele is also shown. (B) Comparison of number of eggs laid by wild type (WT) and *seip-1(-)* animals. At least 10 animals of each genotype were scored. (C) The percentage of BODIPY-stained embryos produced by individual *seip-1(-)* adult during the egg-laying period. 20 animals were monitored. (D) The percentage of BODIPY-stained embryos produced by animals of the indicated genotype, quantified in a defined time window. Five independent biological samples were scored, each with progeny from four 1-day-old adults. For detailed experimental setup, refer to Methods and Materials.

**Figure S2.**
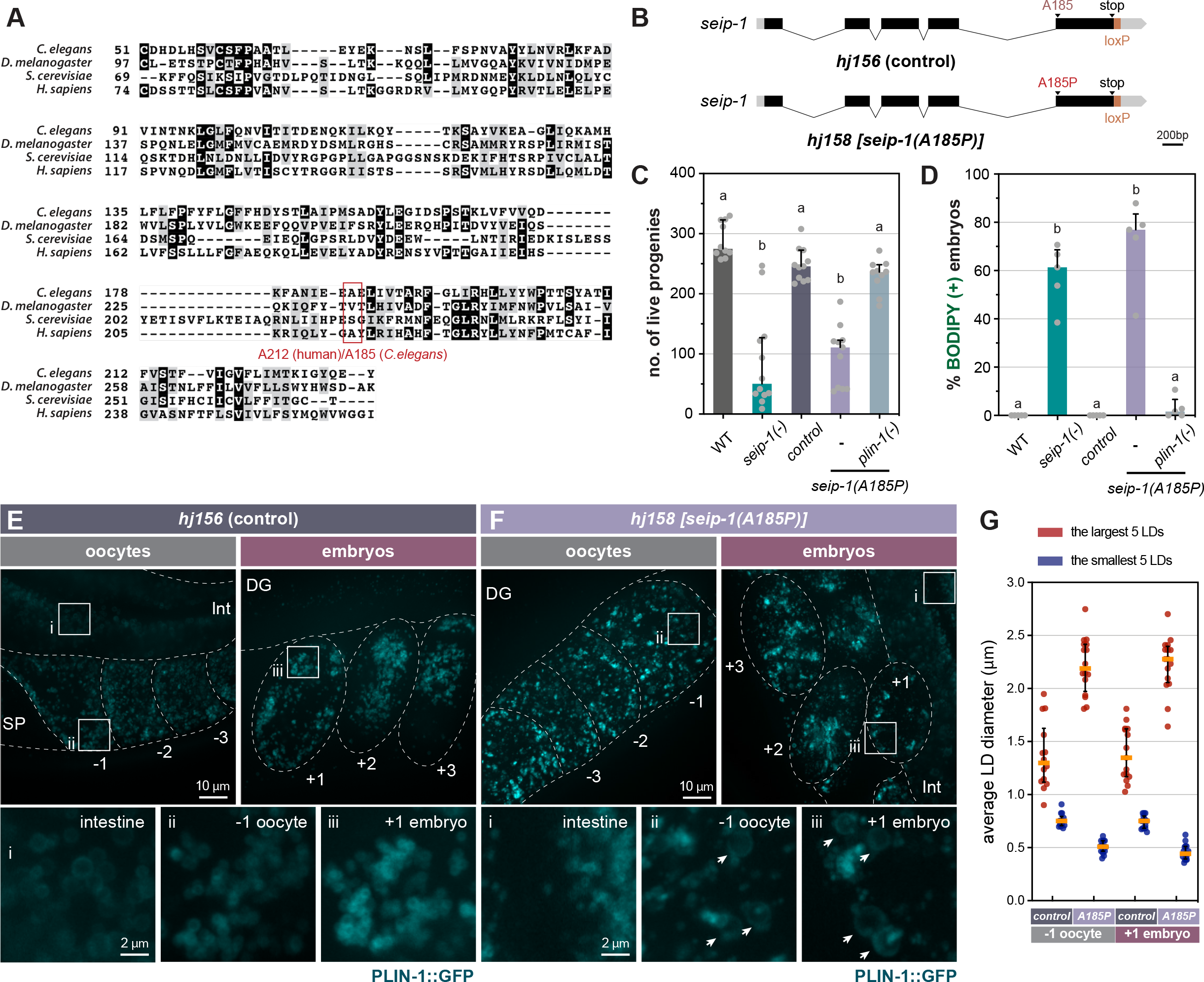
Animals that express SEIP-1(A185P) phenocopy *seip-1(-)* mutants. (A) Sequence alignment of *C. elegans, Drosophila, S. cerevisiae*, and human seipin orthologs.The human lipodystrophy-associated A212P mutation in seipin is equivalent to A185P in *C. elegans* SEIP-1. (B) Schematic representation of specific mutations introduced to the endogenous *seip-1* locus. The two alleles, *hj156* and *hj158*, were engineered by CRISPR/Cas9-mediated HDR with repair templates that only differ at the coding sequence of A185. A loxP scar was left between the stop codon and 3’-UTR of *seip-1* as a result of removing the self-excising cassette (SEC). (C) The total number of live progeny from individual animals. At least 10 animals of each genotype were scored. Groups with different letters are significantly different (ordinary one-way ANOVA with Turkey’s multiple comparisons test, p<0.01). (D) As in (C), but with the percentage of BODIPY-stained embryos quantified in a defined time window. Five independent repeats were scored, each with progeny from four 1-day-old adults. (E) Visualization of LDs using PLIN-1::GFP (*hj178*) in 1-day-old *seip-1(hj156)* (control) adults. mCherry::PH(PLC1d1) (*itIs44*) labels PM in the germline. mCherry and GFP are pseudocolored magenta and cyan, respectively. Dotted lines mark the boundary between different tissues or embryos. Boxed regions were magnified 5x and displayed at the bottom. A projection of 4.5μm z stack reconstituted from 10 focal planes is shown. For anatomical positions of the ROI, refer to Figure 2A. (F) As in (E), but with *seip-1(hj158)* (A185P). Arrows point to aberrantly enlarged LDs. (G) Average diameter of the largest (crimson) or smallest (navy-blue) five LDs in individual 1-day-old adults. At least 10 animals of each genotype were scored. In both oocytes and embryos, when compared to control, the difference between crimson and navy-blue dots is further augmented in *seip-1(A185P)*.

**Figure S3.**
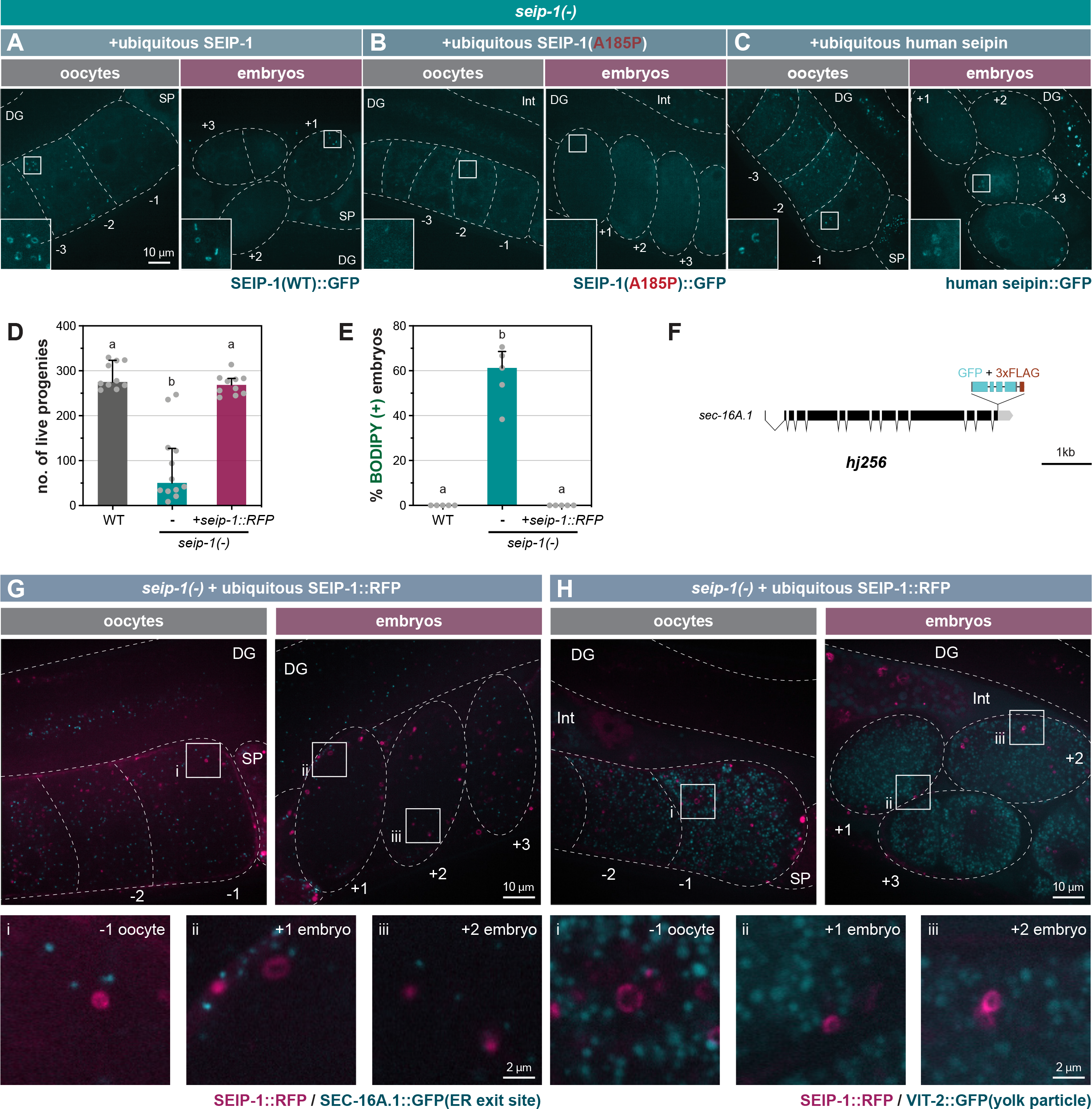
Localization of SEIP-1 in the germline. (A) Visualization of transgenic wild-type SEIP-1::GFP (*hjSi189*) in 1-day-old *seip-1(-)* adults.GFP is pseudocolored cyan. Dotted lines mark the boundary between different tissues or embryos. Boxed regions were magnified 3x and displayed in the inset. A projection of 4.5μm z stack reconstituted from 10 focal planes is shown. For anatomical positions of the ROI, refer to Figure 2A. (B) As in (A), but with transgenic SEIP-1(A185P)::GFP (*hjSi541*). (C) As in (A), but with transgenic human seipin::GFP (*hjSi223*). (D) The total number of live progeny from individual animals. At least 10 animals of each genotype were scored. Groups with different letters are significantly different (ordinary one-way ANOVA with Turkey’s multiple comparisons test, p<0.01). Transgenic SEIP-1::tagRFP is expressed from the single-copy transgene *hjSi434*. (E) As in (D), but with the percentage of BODIPY-stained embryos quantified in a defined time window. Five independent repeats were scored, each with progeny from four 1-day-old adults. (F) Schematic representation of endogenously tagged *sec-16A*.*1 (hj256)*. All isoforms of SEC-16A.1 are fused with C-terminal GFP in *hj256*. (G) Visualization of SEIP-1::tagRFP (*hjSi434*) and SEC-16A.1::GFP (*hj256*) in a 1-day-old *seip-1(-)* adult. tagRFP and GFP are pseudocolored magenta and cyan, respectively. Single focal planes are shown. Dotted lines mark the boundary between different tissues or embryos. Boxed regions were magnified 5x and displayed at the bottom. (H) As in (G), but with an animal that expressed SEIP-1::tagRFP and VIT-2::GFP (*crg9070*).

**Figure S4.**
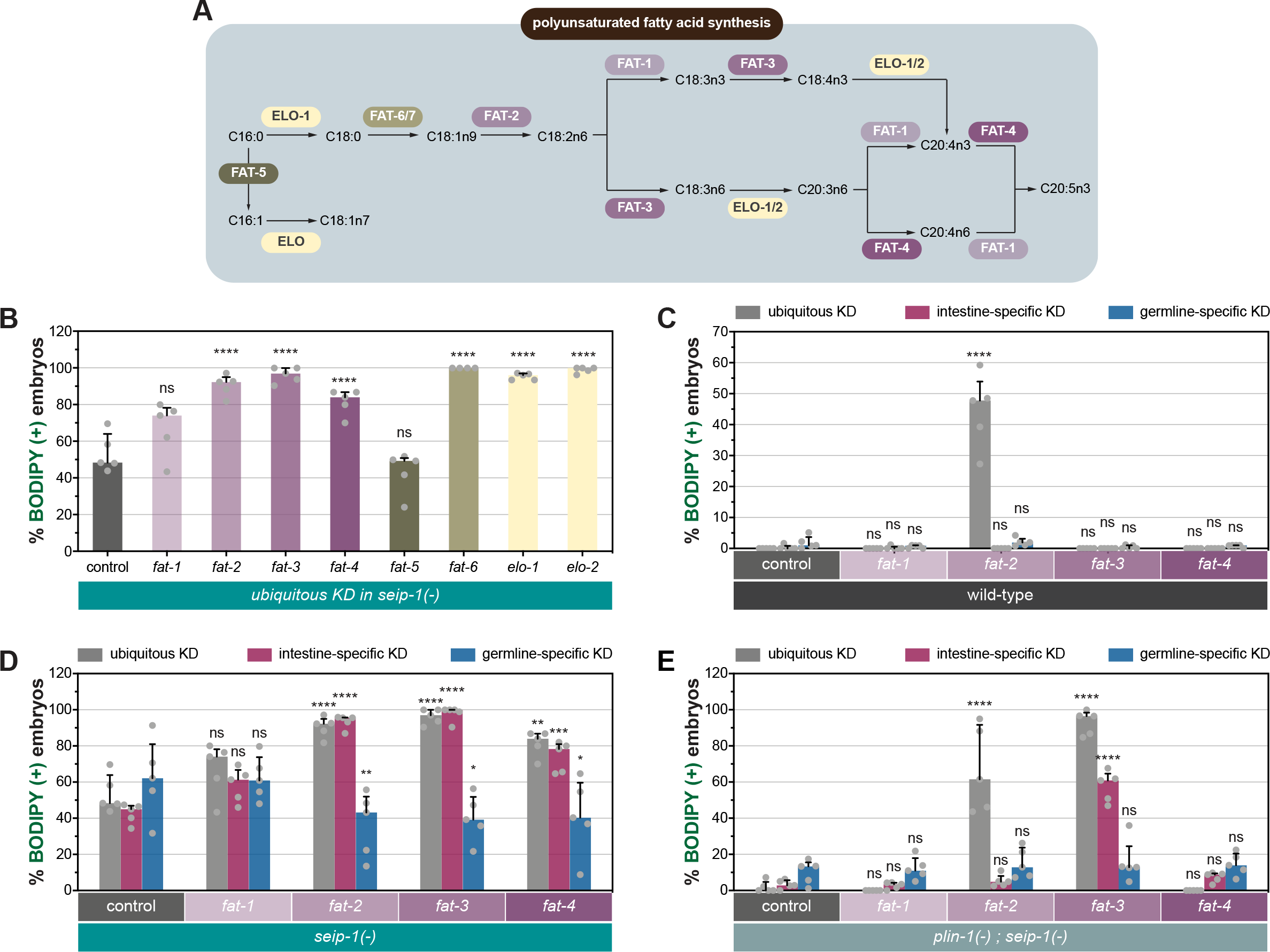
Somatic fatty acid desaturases contribute to eggshell integrity. (A) The biosynthetic pathway of polyunsaturated fatty acids (PUFAs) in *C. elegans*. (B) The percentage of BODIPY-stained embryos in a defined time window upon constitutive depletion of each gene. Five independent replicates were performed, each with progeny from four 1-day-old *seip-1(-)* adults. Statistical significance on top of each bar is calculated by comparing with the control group (one-way ANOVA with Dunnett’s multiple comparisons test). ns, not significant; ****p < 0.0001. (C) As in (B), but with individual genes knocked down in the specified tissues of one-day-old wild-type adults. Statistical significance on top of each bar was calculated by comparing each experimental group with its counterpart in the control group (RNAi vector) (two-way ANOVA and Sidak’s multiple comparisons test). (D) As in (C), but with *seip-1(-)* adults. (E) As in (D), but with *plin-1(-); seip-1(-)* adults.

**Figure S5.**
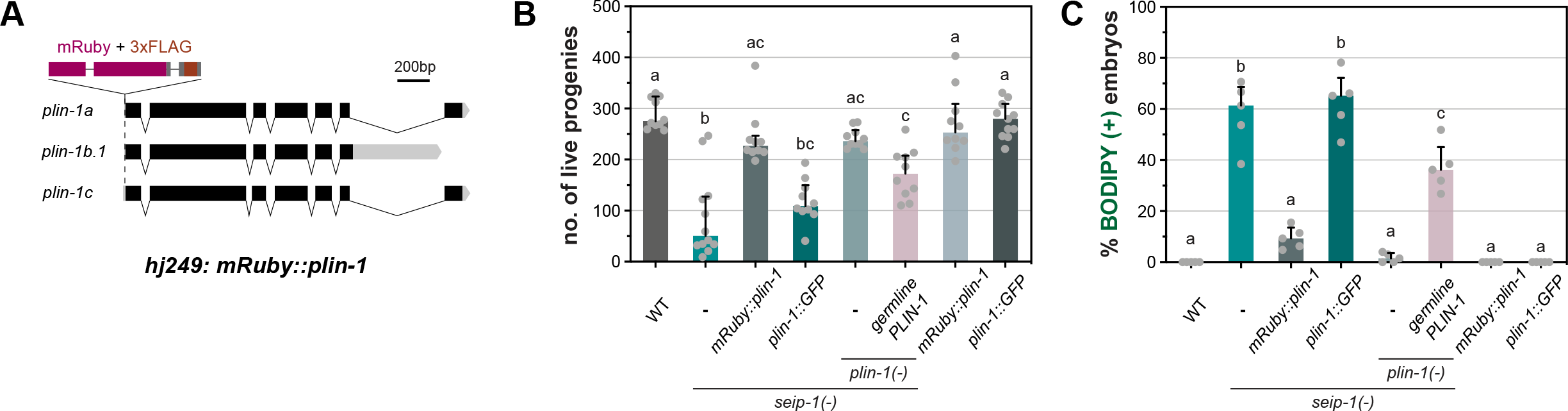
The mRuby::PLIN-1 fusion protein is not fully functional. (A) Schematic representation of the modified *plin-1* locus by CRISPR/Cas9. All isoforms of PLIN-1 are fused with mRuby at the N-terminus in animals that carry the *plin-1(hj249)* allele. (B) The total number of live progeny from individual animals. At least 10 animals of each genotype were scored. Groups with different letters are significantly different (ordinary one-way ANOVA with Turkey’s multiple comparisons test, p<0.01). Germline rescue of *plin-1(-)* was performed by expressing PLIN-1::GFP from a single-copy transgene, driven by the *sun-1* promoter (*hjSi552*). (C) As in (B), but with the percentage of BODIPY-stained embryos quantified in a defined time window. Five independent repeats were scored, each with progeny from four 1-day-old adults.

**Figure S6.**
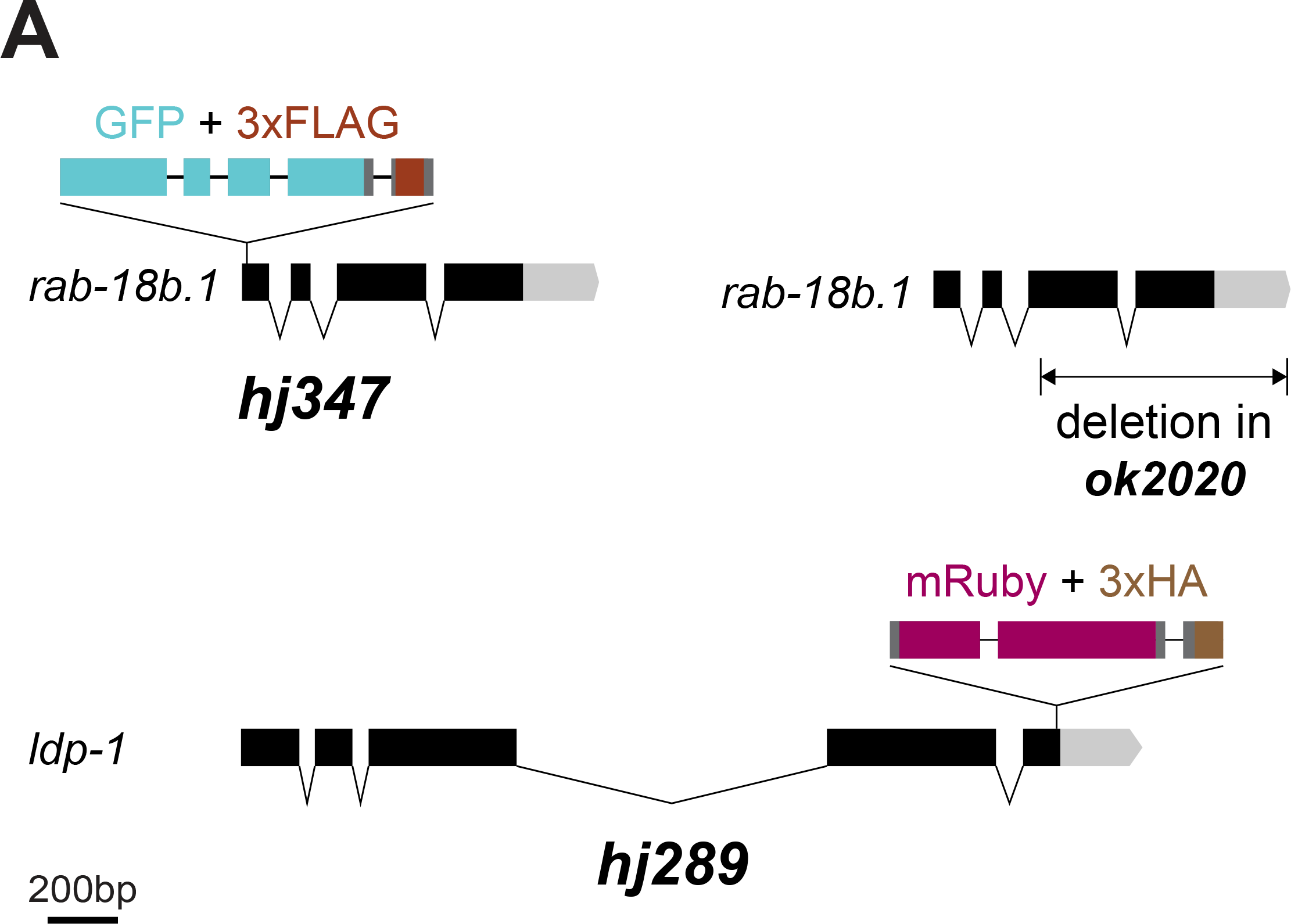
Schematic representation of knock-in or mutant alleles used in Figure 4.

## References

Bai, X., Huang, L.-J., Chen, S.-W., Nebenfuehr, B., Wysolmerski, B., Wu, J.-C., et al. (2020). Loss of the seipin gene perturbs eggshell formation in Caenorhabditiselegans. Dev. Camb. Engl. 147. doi:10.1242/dev.192997.

Bembenek, J. N., Richie, C. T., Squirrell, J. M., Campbell, J. M., Eliceiri, K. W., Poteryaev, D., et al. (2007). Cortical granule exocytosis in C. elegans is regulated by cell cycle components including separase. Development 134, 3837–3848. doi:10.1242/dev.011361.

Binns, D., Lee, S., Hilton, C. L., Jiang, Q.-X., and Goodman, J. M. (2010). Seipin is a discrete homooligomer. Biochemistry 49, 10747–10755. doi:10.1021/bi1013003.

Cao, Z., Hao, Y., Fung, C. W., Lee, Y. Y., Wang, P., Li, X., et al. (2019). Dietary fatty acids promote lipid droplet diversity through seipin enrichment in an ER subdomain. Nat. Commun. 10, 2902. doi:10.1038/s41467-019-10835-4.

Cao, Z., Wang, X., Huang, X., and Mak, H. Y. (2021). Are endoplasmic reticulum subdomains shaped by asymmetric distribution of phospholipids? Evidence from a C. elegans model system. BioEssays News Rev. Mol. Cell. Dev. Biol. 43, e2000199. doi:10.1002/bies.202000199.

Cartwright, B. R., and Goodman, J. M. (2012). Seipin: from human disease to molecular mechanism. J. Lipid Res. 53, 1042–1055. doi:10.1194/jlr.R023754.

Carvalho, A., Olson, S. K., Gutierrez, E., Zhang, K., Noble, L. B., Zanin, E., et al. (2011). Acute drug treatment in the early C. elegans embryo. PloS One 6, e24656. doi:10.1371/journal.pone.0024656.

Chen, W., Chang, B., Saha, P., Hartig, S. M., Li, L., Reddy, V. T., et al. (2012). Berardinelli-Seip Congenital Lipodystrophy 2/Seipin Is a Cell-Autonomous Regulator of Lipolysis Essential for Adipocyte Differentiation. Mol. Cell. Biol. 32, 1099–1111. doi:10.1128/MCB.06465-11.

Chen, W.-W., Lemieux, G. A., Camp, C. H., Chang, T.-C., Ashrafi, K., and Cicerone, M. T. (2020). Spectroscopic coherent Raman imaging of Caenorhabditis elegans reveals lipid particle diversity. Nat. Chem. Biol. 16, 1087–1095. doi:10.1038/s41589-020-0565-2.

Chung, J., Wu, X., Lambert, T. J., Lai, Z. W., Walther, T. C., and Farese, R. V. (2019). LDAF1 and Seipin Form a Lipid Droplet Assembly Complex. Dev. Cell 51, 551-563.e7. doi:10.1016/j.devcel.2019.10.006.

Dickinson, D. J., Pani, A. M., Heppert, J. K., Higgins, C. D., and Goldstein, B. (2015). Streamlined Genome Engineering with a Self-Excising Drug Selection Cassette. Genetics 200, 1035–1049. doi:10.1534/genetics.115.178335.

El Zowalaty, A. E., Li, R., Chen, W., and Ye, X. (2018). Seipin deficiency leads to increased endoplasmic reticulum stress and apoptosis in mammary gland alveolar epithelial cells during lactation. Biol. Reprod. 98, 570–578. doi:10.1093/biolre/iox169.

Ezcurra, M., Benedetto, A., Sornda, T., Gilliat, A. F., Au, C., Zhang, Q., et al. (2018). C. elegans Eats Its Own Intestine to Make Yolk Leading to Multiple Senescent Pathologies. Curr. Biol. 28, 2544-2556.e5. doi:10.1016/j.cub.2018.06.035.

Fei, W., Shui, G., Gaeta, B., Du, X., Kuerschner, L., Li, P., et al. (2008). Fld1p, a functional homologue of human seipin, regulates the size of lipid droplets in yeast. J. Cell Biol. 180, 473–482. doi:10.1083/jcb.200711136.

Fu, D., Yu, Y., Folick, A., Currie, E., Farese, R. V., Tsai, T.-H., et al. (2014). In vivo metabolic fingerprinting of neutral lipids with hyperspectral stimulated Raman scattering microscopy. J. Am. Chem. Soc. 136, 8820–8828. doi:10.1021/ja504199s.

Goldstein, B., Hird, S. N., and White, J. G. (1993). Cell polarity in early C. elegans development. Dev. Camb. Engl. Suppl., 279–287.

Grant, B., and Hirsh, D. (1999). Receptor-mediated Endocytosis in the Caenorhabditis elegans Oocyte. Mol. Biol. Cell 10, 4311–4326.

Green, R. A., Audhya, A., Pozniakovsky, A., Dammermann, A., Pemble, H., Monen, J., et al. (2008). Expression and imaging of fluorescent proteins in the C. elegans gonad and early embryo. Methods Cell Biol. 85, 179–218. doi:10.1016/S0091-679X(08)85009-1.

Habuchi, S., Tsutsui, H., Kochaniak, A. B., Miyawaki, A., and Oijen, A. M. van (2008). mKikGR, a Monomeric Photoswitchable Fluorescent Protein. PLOS ONE 3, e3944. doi:10.1371/journal.pone.0003944.

Huelgas-Morales, G., and Greenstein, D. (2018). Control of oocyte meiotic maturation in C. elegans. Semin. Cell Dev. Biol. 84, 90–99. doi:10.1016/j.semcdb.2017.12.005.

Ibayashi, M., Aizawa, R., Mitsui, J., and Tsukamoto, S. (2021). Homeostatic regulation of lipid droplet content in mammalian oocytes and embryos. Reprod. Camb. Engl. 162, R99–R109. doi:10.1530/REP-21-0238.

Johnston, W. L., and Dennis, J. W. (2012). The eggshell in the C. elegans oocyte-to-embryo transition. Genes. N. Y. N 2000 50, 333–349. doi:10.1002/dvg.20823.

Magré, J., Delépine, M., Khallouf, E., Gedde-Dahl, T., Van Maldergem, L., Sobel, E., et al. (2001). Identification of the gene altered in Berardinelli-Seip congenital lipodystrophy on chromosome 11q13. Nat. Genet. 28, 365–370. doi:10.1038/ng585.

Masedunskas, A., Chen, Y., Stussman, R., Weigert, R., and Mather, I. H. (2017). Kinetics of milk lipid droplet transport, growth, and secretion revealed by intravital imaging: lipid droplet release is intermittently stimulated by oxytocin. Mol. Biol. Cell 28, 935–946. doi:10.1091/mbc.E16-11-0776.

Mitani, S. (2017). Comprehensive functional genomics using Caenorhabditis elegans as a model organism. Proc. Jpn. Acad. Ser. B Phys. Biol. Sci. 93, 561–577. doi:10.2183/pjab.93.036.

Molenaar, M. R., Yadav, K. K., Toulmay, A., Wassenaar, T. A., Mari, M. C., Caillon, L., et al. (2021). Retinyl esters form lipid droplets independently of triacylglycerol and seipin. J. Cell Biol. 220, e202011071. doi:10.1083/jcb.202011071.

Mosquera, J. V., Bacher, M. C., and Priess, J. R. (2021). Nuclear lipid droplets and nuclear damage in Caenorhabditis elegans. PLoS Genet. 17, e1009602. doi:10.1371/journal.pgen.1009602.

Na, H., Zhang, P., Chen, Y., Zhu, X., Liu, Y., Liu, Y., et al. (2015). Identification of lipid droplet structure-like/resident proteins in Caenorhabditis elegans. Biochim. Biophys. Acta 1853, 2481–2491. doi:10.1016/j.bbamcr.2015.05.020.

Neve, I. A. A., Sowa, J. N., Lin, C.-C. J., Sivaramakrishnan, P., Herman, C., Ye, Y., et al. (2020). Escherichia coli Metabolite Profiling Leads to the Development of an RNA Interference Strain for Caenorhabditis elegans. G3 Bethesda Md 10, 189–198. doi:10.1534/g3.119.400741.

Olson, S. K., Greenan, G., Desai, A., Müller-Reichert, T., and Oegema, K. (2012). Hierarchical assembly of the eggshell and permeability barrier in C. elegans. J Cell Biol 198, 731–748. doi:10.1083/jcb.201206008.

Olzmann, J. A., and Carvalho, P. (2019). Dynamics and functions of lipid droplets. Nat. Rev. Mol. Cell Biol. 20, 137–155. doi:10.1038/s41580-018-0085-z.

Ozeki, S., Cheng, J., Tauchi-Sato, K., Hatano, N., Taniguchi, H., and Fujimoto, T. (2005). Rab18 localizes to lipid droplets and induces their close apposition to the endoplasmic reticulum-derived membrane. J. Cell Sci. 118, 2601–2611. doi:10.1242/jcs.02401.

Payne, V. A., Grimsey, N., Tuthill, A., Virtue, S., Gray, S. L., Dalla Nora, E., et al. (2008). The human lipodystrophy gene BSCL2/seipin may be essential for normal adipocyte differentiation. Diabetes 57, 2055–2060. doi:10.2337/db08-0184.

Perez, M. F., Francesconi, M., Hidalgo-Carcedo, C., and Lehner, B. (2017). Maternal age generates phenotypic variation in C. elegans. Nature 552, 106–109. doi:10.1038/nature25012.

Prasanna, X., Salo, V. T., Li, S., Ven, K., Vihinen, H., Jokitalo, E., et al. (2021). Seipin traps triacylglycerols to facilitate their nanoscale clustering in the endoplasmic reticulum membrane. PLoS Biol. 19, e3000998. doi:10.1371/journal.pbio.3000998.

Salo, V. T., Belevich, I., Li, S., Karhinen, L., Vihinen, H., Vigouroux, C., et al. (2016). Seipin regulates ER-lipid droplet contacts and cargo delivery. EMBO J. 35, 2699–2716. doi:10.15252/embj.201695170.

Salo, V. T., Li, S., Vihinen, H., Hölttä-Vuori, M., Szkalisity, A., Horvath, P., et al. (2019). Seipin Facilitates Triglyceride Flow to Lipid Droplet and Counteracts Droplet Ripening via Endoplasmic Reticulum Contact. Dev. Cell 50, 478-493.e9. doi:10.1016/j.devcel.2019.05.016.

Sharrock, W. J., Sutherlin, M. E., Leske, K., Cheng, T. K., and Kim, T. Y. (1990). Two distinct yolk lipoprotein complexes from Caenorhabditis elegans. J. Biol. Chem. 265, 14422–14431.

Stein, K. K., and Golden, A. (2018). The C. elegans eggshell. WormBook Online Rev. C Elegans Biol. 2018, 1–36. doi:10.1895/wormbook.1.179.1.

Sui, X., Arlt, H., Brock, K. P., Lai, Z. W., DiMaio, F., Marks, D. S., et al. (2018). Cryo–electron microscopy structure of the lipid droplet–formation protein seipin. J. Cell Biol. 217, 4080–4091. doi:10.1083/jcb.201809067.

Szymanski, K. M., Binns, D., Bartz, R., Grishin, N. V., Li, W.-P., Agarwal, A. K., et al. (2007). The lipodystrophy protein seipin is found at endoplasmic reticulum lipid droplet junctions and is important for droplet morphology. Proc. Natl. Acad. Sci. 104, 20890–20895. doi:10.1073/pnas.0704154104.

Tagawa, A., Rappleye, C. A., and Aroian, R. V. (2001). Pod-2, along with pod-1, defines a new class of genes required for polarity in the early Caenorhabditis elegans embryo. Dev. Biol. 233, 412–424. doi:10.1006/dbio.2001.0234.

Tauchi-Sato, K., Ozeki, S., Houjou, T., Taguchi, R., and Fujimoto, T. (2002). The surface of lipid droplets is a phospholipid monolayer with a unique Fatty Acid composition. J. Biol. Chem. 277, 44507–44512. doi:10.1074/jbc.M207712200.

Thul, P. J., Tschapalda, K., Kolkhof, P., Thiam, A. R., Oberer, M., and Beller, M. (2017). Targeting of the Drosophila protein CG2254/Ldsdh1 to a subset of lipid droplets. J. Cell Sci. 130, 3141–3157. doi:10.1242/jcs.199661.

Walther, T. C., Chung, J., and Farese, R. V. (2017). Lipid Droplet Biogenesis. Annu. Rev. Cell Dev. Biol. 33, 491–510. doi:10.1146/annurev-cellbio-100616-060608.

Wang, H., Becuwe, M., Housden, B. E., Chitraju, C., Porras, A. J., Graham, M. M., et al. (2016). Seipin is required for converting nascent to mature lipid droplets. eLife 5, e16582. doi:10.7554/eLife.16582.

Watson, P., Townley, A. K., Koka, P., Palmer, K. J., and Stephens, D. J. (2006). Sec16 Defines Endoplasmic Reticulum Exit Sites and is Required for Secretory Cargo Export in Mammalian Cells. Traffic Cph. Den. 7, 1678–1687. doi:10.1111/j.1600-0854.2006.00493.x.

Welte, M. A. (2015). As the fat flies: The dynamic lipid droplets of Drosophila embryos. Biochim. Biophys. Acta 1851, 1156–1185. doi:10.1016/j.bbalip.2015.04.002.

Wilfling, F., Wang, H., Haas, J. T., Krahmer, N., Gould, T. J., Uchida, A., et al. (2013). Triacylglycerol synthesis enzymes mediate lipid droplet growth by relocalizing from the ER to lipid droplets. Dev. Cell 24, 384–399. doi:10.1016/j.devcel.2013.01.013.

Xie, K., Zhang, P., Na, H., Liu, Y., Zhang, H., and Liu, P. (2019). MDT-28/PLIN-1 mediates lipid droplet-microtubule interaction via DLC-1 in Caenorhabditis elegans. Sci. Rep. 9, 14902. doi:10.1038/s41598-019-51399-z.

Xu, N., Zhang, S. O., Cole, R. A., McKinney, S. A., Guo, F., Haas, J. T., et al. (2012). The FATP1–DGAT2 complex facilitates lipid droplet expansion at the ER–lipid droplet interface. J. Cell Biol. 198, 895–911. doi:10.1083/jcb.201201139.

Zoni, V., Khaddaj, R., Lukmantara, I., Shinoda, W., Yang, H., Schneiter, R., et al. (2021). Seipin accumulates and traps diacylglycerols and triglycerides in its ring-like structure. Proc. Natl. Acad. Sci. U. S. A. 118, e2017205118. doi:10.1073/pnas.2017205118.

Zou, L., Wu, D., Zang, X., Wang, Z., Wu, Z., and Chen, D. (2019). Construction of a germline-specific RNAi tool in C. elegans. Sci. Rep. 9, 2354. doi:10.1038/s41598-019-38950-8.

